# Deep proteome profiling with reduced carry over using superficially porous microfabricated nanoLC columns

**DOI:** 10.1101/2021.11.28.470272

**Authors:** Karel Stejskal, Jeff Op de Beeck, Manuel Matzinger, Gerhard Dürnberger, Alexander Boychenko, Paul Jacobs, Karl Mechtler

**Affiliations:** IMP - Institute of Molecular Pathology, Campus-Vienna-Biocenter 1, A-1030 Vienna, Austria; IMBA - Institute of Molecular Biotechnology of the Austrian Academy of Sciences, Dr. Bohr Gasse 3, A-1030 Vienna, Austria; Gregor Mendel Institute of Molecular Plant Biology of the Austrian Academy of Sciences, Dr. Bohr Gasse 3, A-1030 Vienna, Austria; Thermo Fisher Scientific, Technologiepark-Zwijnaarde 82, B-9052 Gent, Belgium; Thermo Fisher Scientific, Dornierstrasse 4, 82110 Germering, Germany

## Abstract

In the field of LC-MS based proteomics, increases in sampling depth and proteome coverage have mainly been accomplished by rapid advances in mass spectrometer technology. The comprehensiveness and quality of data that can be generated do however also depend on the performance provided by nano liquid chromatography (nanoLC) separations. Proper selection of reversed-phase separation columns can be important to provide the MS instrument with peptides at the highest possible concentration and separated at the highest possible resolution. In the current contribution, we evaluate the use of prototype generation 2 μPAC nanoLC columns which use C18 functionalized superficially porous micro pillars as a stationary phase. When comparing to traditionally used fully porous silica stationary phases, more precursors could be characterized when performing single shot data-dependent LC-MS/MS analyses of a human cell line tryptic digest. Up to 30% more protein groups and 60% more unique peptides were identified for short gradients (10 min) and limited sample amounts (10-100 ng of cell lysate digest). With LC-MS gradient times of 10, 60, 120 and 180 min, we respectively identified 2252, 6513, 7382 and 8174 protein groups with 25, 500, 1000 and 2000 ng of sample loaded on column. Reduction of sample carry over to the next run (up to 2 to 3%) and decreased levels of methionine oxidation (up to 3-fold) were identified as additional figures of merit. When analyzing a disuccinimidyl dibutyric urea (DSBU) crosslinked synthetic library, 29 to 59 more unique crosslinked peptides could be identified at a experimentally validated false discovery rate (FDR) of 1-2%.

## INTRODUCTION

Even though the practice of LC-MS based bottom-up proteomics has remained relatively unaltered over the past decade, researchers are progressively closing the gap between experimentally identified and theoretically expected proteoforms present in complex cell lysates^1–3^. Key aspects driving this progress are the continuous evolution of MS/MS instruments, the coming of age of additional ion mobility separation techniques and the combination with LC separation that delivers maximal resolving power and throughput^1,4,5^. Even though MS/MS instruments have evolved to a point where acquisition rates up to 133 Hz can be reached^6,7^, these developments have struggled to materialize similar leaps in proteome coverage depth such as obtained by publications of Thakur et al., Hebert et al. and Scheltema et al in 2011 and 2014^8–10^. As postulated several years ago by Shishkova et al^11^, chromatographic separation performance is the key but perhaps underappreciated bottleneck limiting the speed and depth of single-shot proteomic analyses. Improvements in a chromatographic resolution have historically been achieved by increasing column length or by decreasing silica particle diameters^12–14^. However, reducing particle diameters and extending column length have a synergistic effect on operating pressures^15,16^. Consequently, current state-of-the art nano LC columns often require ultra-high pressure liquid chromatography (UHPLC) instruments which can accurately deliver nanoliter per minute flow rates at operating pressures up to 1,500 bar.

To cope with these pressure requirements, alternative formats such as monolithic columns have been introduced, albeit with limited adoption in the field of proteomics^17–19^. Alternatively, microfabricated pillar array columns (μPAC) have been proposed as a new promising technology that can redefine the boundaries of LC performance^20^. Using micromachining techniques rather than slurry packing, both chromatographic performance and column permeability can be controlled by design. We describe the use of a new generation of pillar array columns where design specifications have been tightened in search of increased separation performance. Schematic drawings of the ‘building’ blocks or unit cells used to design different ‘generations’ of pillar array columns are shown in figure S1. Analogous to observations in packed bed columns, reduction of the pillar and inter pillar dimensions by a factor of 2 results in a net gain in separation resolution with a factor of 1.4 at the cost of increased operating pressure^21^. In contrast to the experiments we conducted for limited sample amounts in 2021^22^, the work we report on in the current contribution uses a superficially porous rather than a non-porous version of the generation 2 pillar array column. By using electrochemical anodization, the outer shell of the cylindrical pillars is rendered mesoporous with pore sizes in the range of 100 to 300 Å. This increases the available interaction surface by a factor of aproximately 30, making this format more compatible with conventional sample loads.

To investigate potential benefits of the μPAC column for nanoLC-MS applications, we report on an extensive benchmarking series where we coupled this column to the latest generation of tribrid MS systems, a Field Asymmetric Waveform Ion Mobility Spectrometry (FAIMS) pro interface and a next-generation low-flow UHPLC system (Vanquish Neo UHPLC). Such experiments are commonly performed with highly validated mammalian protein digest standards to provide unbiased data on instrument performance. Results do however often differ from what can be achieved with biologically relevant samples and fail to provide information on day-to-day robustness and throughput. The current study aims to address these matters by providing additional data on performance over time, column related sample carry over and validation of results by implementing the workflow for the analysis of a synthetic library of cross-linked peptides^23^.

## EXPERIMENTAL

### Sample preparation

Pierce™ HeLa Protein Digest Standard (ThermoFisher Scientific) was used for MS parameter optimization as well as for final column benchmarking measurements. 20 μg peptide pellets were dissolved in LC/MS grade water with 0.1% (v/v) TFA and diluted to the required peptide concentration in autosampler vials (Fisherbrand™ 9 mm Short Thread TPX Vial with integrated Glass Micro-Insert; Cat. No. 11515924). All liquid handling was done as fast as possible without unnecessary time gaps with the aim to minimize sample losses on plastics and glass surfaces.

For the cross-linking experiment, synthetic peptides generated by Beveridge and coworkers were cross-linked using DSBU exactly as described in their paper^23^. The final cross-linked peptide mix was merged either with an equal amount of tryptic HeLa peptides (Pierce™ HeLa Protein Digest Standard dissolved in 0.1% TFA) to obtain 1:1 spiked system or with the 5x mass of tryptic HeLa peptides to obtain a 1:5 spiked system. A total amount of 1 μg (either using the cross-linked peptide-mix only or total peptide after spiking) peptide was used for each LC-MS/MS analysis.

### Liquid chromatography-mass spectrometry analysis (LC-MS)

Peptide samples were analyzed using a Vanquish™ Neo UHPLC instrument in nano/cap mode and configured for direct injection onto the column. Orbitrap Eclipse™ Tribrid™ mass spectrometer equipped with the FAIMS Pro interface (ThermoFisher Scientific). Peptides were separated with either the new generation prototype 50 cm pillar array column (Thermo Fisher) or with a 25 cm length packed bed column with integrated tip.

The 50 cm μPAC was placed in a Butterfly heater (PST-BPH-20, Phoenix S&T) and operated at 50°C. The column was connected to an EASY-Spray™ bullet emitter (10 μm ID, ES993; Thermo Fisher Scientific) with a custom-made fused silica capillary (20 μm ID X 360 μm OD, length 10 cm, Polymicro) with 1/16” ZDV fittings and a precision cut PEEK sleeve on the ESI source facing side. An electrospray voltage of 2.4 kV was applied at the integrated liquid junction of EASY-Spray™ emitter. To avoid electric current from affecting the upstream separation column, a stainless steel 50 μm internal bore reducing union (VICI; C360RU.5S62) was electrically connected to the grounding pin at the pump module (Figure S2).

The packed bed analytical column (25 cm x 75 μm ID, 1.6 μm C18; AUR2-25075C18A; IonOpticks) was installed in a Sonation column oven (PRSO-V2; Sonation) and operated at 50°C. The Sonation column oven was mounted on a NanoFlex™ ion source (ThermoFisher Scientific). An electrospray voltage of 2.4 kV was applied at the nanoZero® fitting via the high voltage cable (HVCABLE01; IonOpticks)(Figure S2).

Peptides were separated with stepped linear solvent gradients, all performed at a flow rate of 200 nL/min (except flow rate experiment) with various durations of 10, 60, 120 and 180 min. Organic modifier content (acetonitrile acidified with 0.1% v/v formic acid) was first increased from 0.8% to 18% in 7.5, 45, 90 and 135 min, then increased from 18% to 32% in, respectively 2.5, 15, 30 and 15 min, and finally ramped from 32% to 76% in 5 min. Mobile phase composition was kept at a high organic phase (76% acetonitrile acidified with 0.1% v/v formic acid) for 5 min to wash the column. Column re-equilibration was performed at a low organic phase (0.8% acetonitrile acidified with 0.1% v/v formic acid) with 2 column volumes.

### MS Acquisition

The mass spectrometer was operated in data-dependent mode, using a full scan with *m/z* range 375-1500, orbitrap resolution of 120.000, target value 250 %, and maximum injection time set to Auto. Compensation voltages of −45, −55 and −75 V or −45, −55, −65 and −75 V were combined in a single run with total cycle times of respectively 3 and 4 s. The intensity threshold for precursor was set to 5e4. Dynamic exclusion duration was based on the length of the LC gradient set up for 10 min to 20 s, for 60 min to 25 s, for 120 min to 40 s and for 180 min to 60 s.

MS/MS spectra were acquired in the ion trap analyzer and fragmented by stepped higher-energy collisional dissociation (HCD) using anormalized collision energy (NCE) of 30 %. Precursors were isolated in a window of 1.0 Da. The linear ion trap acquired spectra in the Turbo mode and in the range 200-1400 m/z. Normalized AGC target was set to 300 % and maximum injection time was 12.5 ms for 10 and 60 min long gradient methods and 15 ms for 120 and 180 min long gradient methods.

### Data analysis

MS/MS spectra from raw data were imported to Proteome Discoverer (PD) (version 2.5.0.400, Thermo Scientific). First, spectra were recalibrated in the PD node “Spectrum Files RC” using the human SwissProt database (Homo sapiens; release 2020_12; 20,541 sequences, 11,395,748 residues) and a database of common contaminants (375 sequences, 144,816 residues). Recalibration was performed for fully tryptic peptides applying an initial precursor mass tolerance of 20 ppm and a fragment mass tolerance of 0.5 Da. Carbamidomethylation of cysteine was set as a fixed modification in the recalibration step. Database search on individual raw files was performed using MS Amanda^24^ (version 2.5.0.16129) and the FASTA databases already described above at recalibration. Trypsin was specified as proteolytic enzyme, cleaving after lysine (K) and arginine (R) except when followed by proline (P) and up to one missed cleavage was considered. Mass tolerance was limited to 7 ppm at the precursor and 0.3 Da at the fragment level. Carbamidomethylation of cysteine (C) was set as a fixed modification and oxidation of methionine (M), as well as acetylation and the loss of methionine at the protein N-terminus were set as a variable modification. Identified spectra were rescored using Percolator ^25^ as implemented in PD and filtered for 1% FDR at the peptide spectrum match and peptide level. Abundance of identified peptides was determined by label free quantification (LFQ) using IMP-apQuant without match beween run mode (MBR) ^26^.

Cross-linked peptides were identified using MS Annika ^27^ (v1.0.18345) within Proteome Discoverer v2.5.0.400. The workflow tree consisted of the MS Annika Detector node (MS tolerance 10 ppm, cross-link modification: DSBU +196.085 Da at lysine, doublet pair selection in combined mode) followed by MS Annika Search (full tryptic digest, 5/10 ppm peptide/fragment mass tolerance, max 3 missed cleavages, carbamidomethyl +57.021 Da at cysteine as static and oxidation +15.995 Da at methionine as dynamic modification) and completed with MS Annika Validator (1 % FDR cutoff at the cross-link specific mathes (CSM) and cross-link (XL) level, separate Intra/Inter-link FDR set to false). Relative abundances of the identified cross-linked peptides were determined by label-free quantification (LFQ) without match between runs using IMP-apQuant^26^. The search was performed against a database containing *S. pyogenes* Cas9 and 116 CRAPome proteins ^28^. For FDR control, peptides cross-linked within the same group (as defined previously) were considered correct, link-connections between peptides of different groups or to peptides from the contaminant database were considered incorrect.

## RESULTS & DISCUSSION

### Column benchmarking

After optimization of a confined set of LC and MS parameters (information provided in supporting information, Figures S3-6 and Tables S1-3), we performed a comprehensive benchmarking experiment to evaluate the column’s applicability for a range of LC gradient settings. The prototype μPAC column was benchmarked against a commercially available packed bed nanoLC column using conditions listed in table S4. When applying a short 10 min gradient (Figure 1A), over 2600 proteins could repeatedly be identified from 100 ng of HeLa digest. A significant increase in both peptide and protein group identifications was observed when comparing the micro pillar array and the packed bed column (student t-test, p<0.001). Even though processed results did not reveal a significant impact on chromatographic performance (peak capacity – Figure S7, median FWHM of peptides – Figure S8,), and the difference in column void times was found to be minimal (6.5 min for the packed bed column and 8.5 min for the μPAC column), 20-30% more protein groups and 40-60% more peptide groups could be identified when using the pillar array format. When plotting the amount of unique PSMs versus retention time (Figure 1B), a clear trend is revealed with additional unique identifications towards the end of the gradient. These data suggest that the column morphology has an impact on the elution behavior of hydrophobic peptide species. Consistent with results reported on the use of superficially porous and large mesopore size stationary phases, we hypothesize that the use of superficially porous rather than fully porous chromatographic media promotes elution and prevents persistent adsorption of analytes to the chromatographic support material. Additional data that confirm this statement are provided when evaluating sample carry over and analyzing cross-linked peptides on both LC column formats. It must be noted that the use of superficially porous stationary phases brings along some limitations, as they typically have lower loading capacity and show poor retention of hydrophilic peptides. Another consideration concerning these fast gradients for low sample amounts is that these methods are far from optimal when maximum instrument occupation efficiency is pursued, as it takes 35 min to have 10 min of peptide elution.

**Figure 1:**
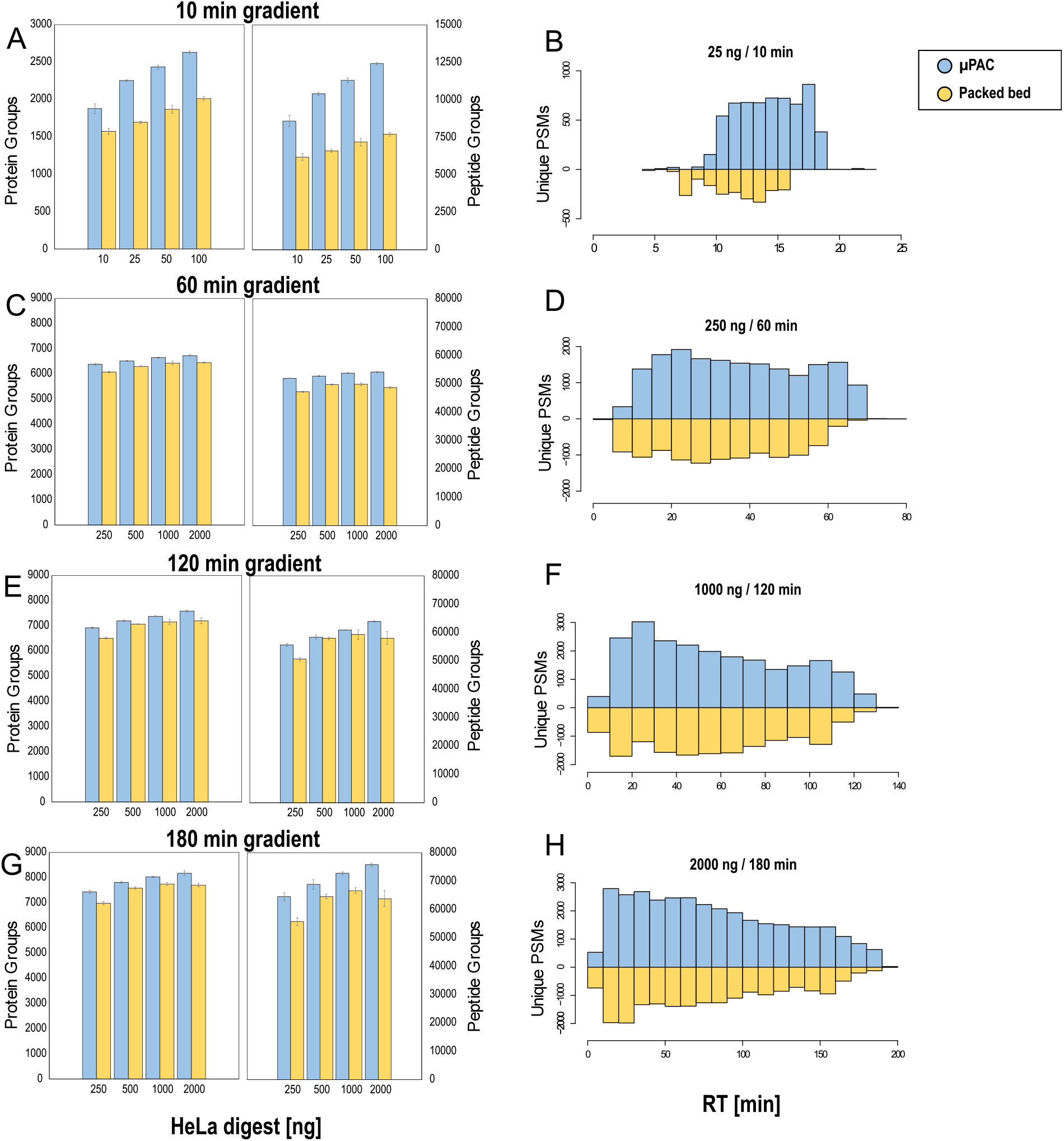
Proteome coverage (protein and peptide Group ID’s) obtained for different gradient lengths and sample loads during the extensive benchmarking experiment. Four different methods are tested, the generation 2 pillar array column (blue) is compared to a packed bed column (Yellow). All values represent average values (n=3, injection replicate) with error bars depicting standard deviations. Unique PSMs identified on each column are plotted as a function of elution time to the right. A-B: 10 min gradient separation, 3 CV FAIMS method, 10-100 ng HeLa digest sample, C-D: 60 min gradient separation, 3 CV FAIMS method, 250-2000 ng HeLa digest sample, E-F: 120 min gradient separation, 3 CV FAIMS method, 250-2000 ng HeLa digest sample, G-H: 180 min gradient separation, 4 CV FAIMS method, 250-2000 ng HeLa digest sample.

More efficient MS utilization can be achieved when using longer solvent gradients (75% for 60, 86% for 120 and 90% for 180 min, MS efficiency calculation provided in supporting information). However, protein identification rates observed for short gradients attenuate according to gradient length. This can be attributed to the fact that the first proteins to be identified from a complex mixture are high abundant ones that can be picked up relatively easy. Further increases in proteomic depth progressively become more challenging as undiscovered proteins are of ever decreasing abundance. This is clearly illustrated in Figure S10, where the abundancies of proteins uniquely discovered by extending gradient length or sample load have been compared to those shared with shorter analyses. The relative increase in protein identifications fades with increasing gradient length, reaching an averaged maximum of close to 8100 protein groups identified out of 2 μg of HeLa digest sample (Figure 1C, E, G). Again, consistently more features were identified when using the pillar array as compared to the packed bed column. Even though the relative increase in identifications was smaller as compared to the high throughput method (3-6% on the protein group level, 6-19% on the peptide group level), unique hits were again predominantly originating from later eluting peptide species (Figure 1D, F, G), confirming earlier observations.

### Artefactual methionine oxidation of peptides

When comparing both column set-ups, a significantly higher portion of peptides containing oxidized methionine residues was identified with the packed bed column (Figure 2B). Even though methionine oxidation of peptides is often biologically relevant (in vivo modification) and for instance observed in a range of oxidative stress and age-related disease states, sample handling and analysis can induce artefactual oxidation (in vitro oxidation) and lead to biased interpretation of biological results. Artefactual oxidation can occur in different stages of a typical bottom-up proteomics workflow, ranging from protein storage and purification to LC separation and ionization^29^. When oxidized species are present within the sample prior reversed phase LC (RPLC) analysis, a retention time difference between the oxidized and the non-oxidized form of the methionine containing peptide is typically observed. The oxidation of a methionine residue to methionine sulfoxide or methionine sulfone reduces hydrophobicity and therefore results in most cases in reduced RPLC retention^30^. LC separation or electrospray ionization induced oxidation on the other hand have much less an impact on peptide retention behavior ^31^. Peptides are not yet present in their oxidized form upon injection onto the LC column and therefore elute much closer to their non-oxidized form. When plotting the amount of oxidized methionine containing peptides as a function of the relative retention time difference with their non-oxidized form (Figure 2A), clear differences are observed between both column setups. Significantly more oxidized features with retention time differences smaller than 2 min are detected when working with the pulled tip emitter column setup (paired t-test between two sample groups from 60, 120 and 180 min gradients, p=0.0127). Up to 54% of oxidized species show retention time differences below 2 minutes, whereas this is only 11% when working with the μPAC column setup.

**Figure 2:**
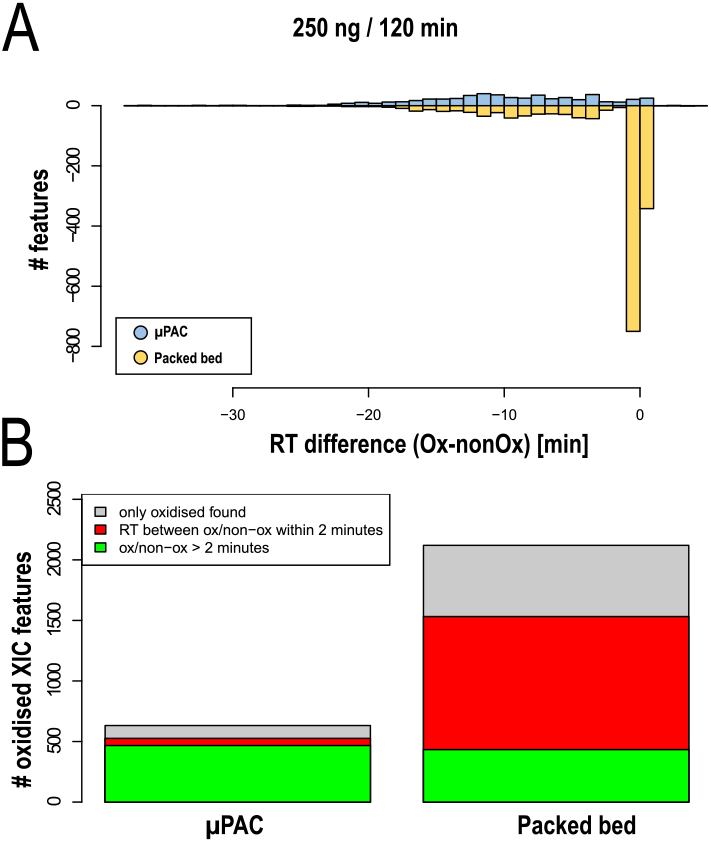
Comparison of peptide oxidation levels between LC column setups. A: number of oxidized methionine containing peptides plotted as a function of the retention time difference between the non-oxidized and the oxidized form – RT difference (ox – nonOx). Generation 2 pillar array column setup (blue) vs packed bed column setup (Yellow). B: Total number of oxidized methionine containing features found for different column setups.

As both columns had very limited operational history (≤ 100 runs, only ‘clean’ digest standards) prior to these analyses and samples were always freshly reconstituted from lyophilized HeLa digest pellets, we suggest that the electrical configuration is the most probable source of on column methionine oxidation. Pulled tip emitter column types require upstream high voltage supply, whereas in contrast, μPAC columns are recommended (or ‘need’) to be plumbed in such a way that a grounded liquid junction shields any effect of the high voltage that is applied downstream at the emitter (Figure S2). The difference observed in oxidized species is consistent with observations described in Liu et al.^32^ hypothesizing on-column oxidation by electrochemically formed radicals when a high potential is applied upstream.

### Carry over

In many cases, very few or no intermediate washes are performed between runs. It is often assumed that a single blank injection is sufficient to clear persistent sample material, without actually acquiring or analyzing MS/MS data. In practice, these assumptions can have a serious impact on results and affect the outcome of a study. The newly introduced Vanquish Neo UHPLC enabled an unbiased investigation of LC column related sample carry over, as after each injection and in parallel to the peptide separation step, the autosampler executes stringent system washing cycles with a high volume of organic solvent to wash the needle outside and the complete injection fluidics path including needle inside. To assess LC column related sample carry over, blank injections were included in the benchmarking series. Figure 3A shows the number of protein groups identified from blank runs for both column set-ups. Up to a sample load of 1 μg, no protein groups were identified from the blank injections on the μPAC prototype, as there were too few spectra present for FDR assessment in Percolator (200 peptides required)^25^. More data is provided by analyzing results obtained for consecutive washes (n=2) that have been performed after 100 ng HeLa QC runs. Using a fixed value validator for FDR assessment, apQuant areas obtained for the top 50 most abundant peptides have been compared (Figure 3C-D). Wash runs performed immediately after the analytical run (1^st^ wash) still show up quite some quantifiable signals for both columns, respectively 43 and 46 out of 50 peptides were quantified on pillar array and packed bed respectively. There is however a significant difference when analyzing data from the 2nd wash run. Respectively 5 and 19 peptides were quantified in the 2^nd^ wash. As mentioned before when discussing the impact of stationary phase support morphology on peptide elution, we believe this is a result of the intrinsic difference in interaction surface between both column types. This consistently results in decreased carry over related identifications on the μPAC column, 3-4 times less on the peptide group and 2-3 times less on the protein group level. Additional experiments with packed bed columns that contain superficially porous particles might provide additional insights to support our statements.

**Figure 3:**
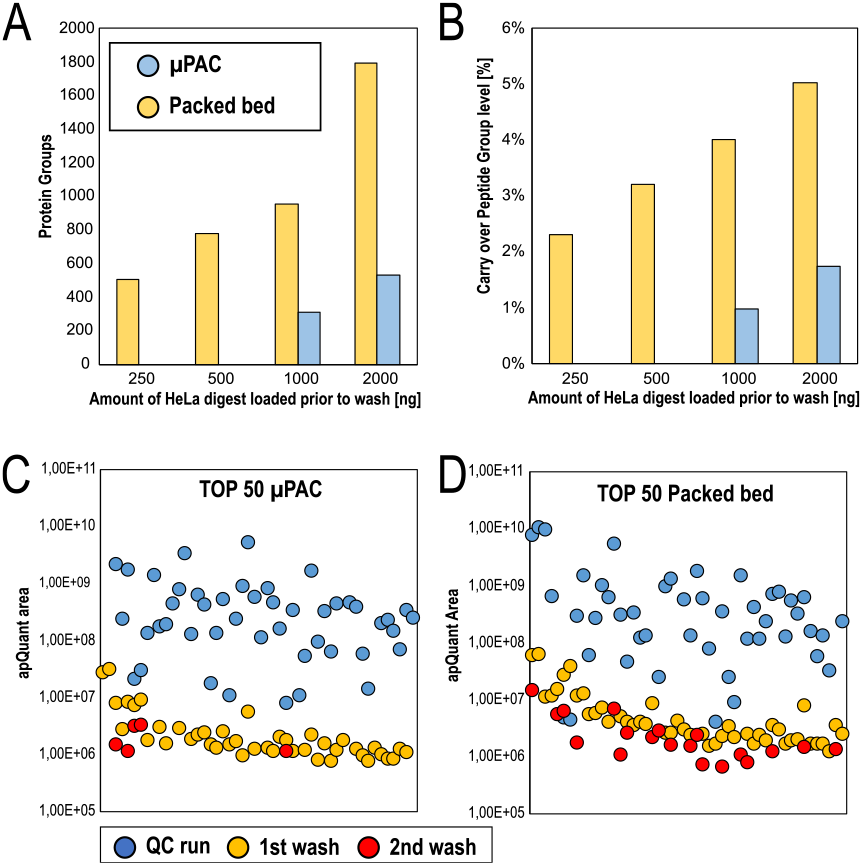
Comparison of sample carry over obtained after increasing sample loads. Blank wash runs immediately after each concentration have been analyzed. Comparison of generation 2 pillar array (blue) to packed bed column (yellow). A: Number of protein group ID’s, B: Relative percentage of carry over on the peptide group level, C-D: Comparison of apQuant area obtained for top 50 most abundant HeLa peptides, 100 ng QC run is compared to results from a first and a second wash, (C = generation 2 pillar array column, D = packed bed column).

### Performance consistency

As from the initial column installation, we implemented a quality control method to assess performance at regular intervals over time. To limit the impact on total acquisition time, a 15 min gradient separation of 100 ng HeLa digest with a total cycle time of 35 min in between runs was used. Consistent performance was obtained throughout a period of 1 month, time which was needed to run MS optimization, LC optimization and actual in-depth benchmarking of a single column. During this period, a slight decrease in protein group ID’s (approximately 14%) was observed, going from 3382 to 2979 protein groups with a single column to emitter assembly. A clear effect is however observed when the pillar array was replaced by a packed bed column, clearly marked by a sudden drop in identifications (Figure S11). Similar proteome depth was never achieved with the packed bed column, resulting in an average of about 2595 protein groups over a 10-day period of analysis time. To confirm that these observations were linked to the LC column type rather than to the MS performance or to batch effects, a second pillar array column was installed immediately after the packed bed column benchmark. This event is again marked by a distinct increase in identifications (Figure S11). Additional experiments to investigate μPAC column-to-column reproducibility were conducted more than a year after the benchmarking experiments. With a similar set-up but now coupled to an Orbitrap Exploris 480 instrument, the performance of 3 prototype μPAC generation 2 columns was compared for 30 min gradient separations. Results have been compiled in the supporting information (Figure S12), showing inter column retention time variation below 1% CV and providing clear proof for consistent performance in bottom-up proteomics with a variation on identifed protein and peptide groups respectively below 1 and 2% CV respectively.

### Cross-linking experiments

In addition to providing a column performance comparison for standardized HeLa digest samples, we performed a limited set of experiments with cross-linked peptide samples. During the last decade, cross-linking mass spectrometry was established as a potent technique to investigate protein-protein interaction networks as well as in the field of structural proteomics. This technique, including a wide variety of applications was already described in several excellent reviews (e.g.^33–35^). Briefly, two amino acid residues are covalently connected by application of the cross-linker reagent, followed by proteomic sample preparation, and yielding two interconnected peptides for detection by mass spectrometry. Depending on the used linker type, cross-linkers can target amines (lysines), sulfhydryl groups (cysteines), carboxylic acids (glutamic- or aspartic-acid) or they can form radical species reactive to any amino acid. The broad variety of linker-types, acquisition techniques as well as data analysis algorithms makes it difficult to find an optimal workflow for a specific protein system. To alleviate this issue, we previously developed a synthetic peptide library based on sequences of the Cas9 protein^23^. The peptides contain exactly one targeted (i.e. lysine) amino acid for cross-linking. They were mixed into groups that were separately cross-linked, followed by quenching and pooling to a single peptide-library. In contrast to experiments where the FDR is computationally determined only by applying the known target-decoy approach, this system allows an exact FDR calculation as only interpeptide connections within a group are possible^36^. Furthermore, the maximal theoretical cross-link number is known (426 unique combinations), which allows estimating the efficiency of a detection workflow based on the reached identification numbers. Such a synthetic library, in combination with the linker reagent (DSBU), therefore represents an ideal benchmarking tool for the comparison of two different chromatographic setups, as done in this study.

The number of identified unique cross-links, as well as the number of cross-link spectrum matches, is reproducibly boosted when using the pillar array column setup compared to the packed bed setup (Figure 4 A, B). The number of identifications decreases upon increasing background of linear peptides present in the sample, which is likely not only a result of increased sample complexity but also of decreased amounts of cross-linked peptides present in spiked samples (i.e. 1μg XL-peptides without spiking vs. 200 ng XL peptides + 800 ng tryptic HeLa peptides in 1:5 spiked sample). Of note, the advantage of the pillar array over packed bed increases in complex sample mixtures. On average, we observed a boost in cross-link IDs of ∼29 % without spiking but of 37% and 59% upon 1:1 and 1:5 spiking respectively. In line with ID numbers also the relative abundance of cross-linked peptides is increased in the pillar array setup for all three test samples. The average real FDR rate is close to the expected 1 % in all sample types and is independent of the used column, highlighting the quality of the obtained data and a properly working target decoy-based FDR approach using MS Annika. As shown in Figure 4C and in line with the results obtained using different sample loads and gradient lengths (Figure 1B, D, F, H), we observed most of the extra CSM identifications with the μPAC column at high retention times. This could indicate fewer losses of larger species which are expected predominant among cross-linked peptides as two peptides are connected.

**Figure 4:**
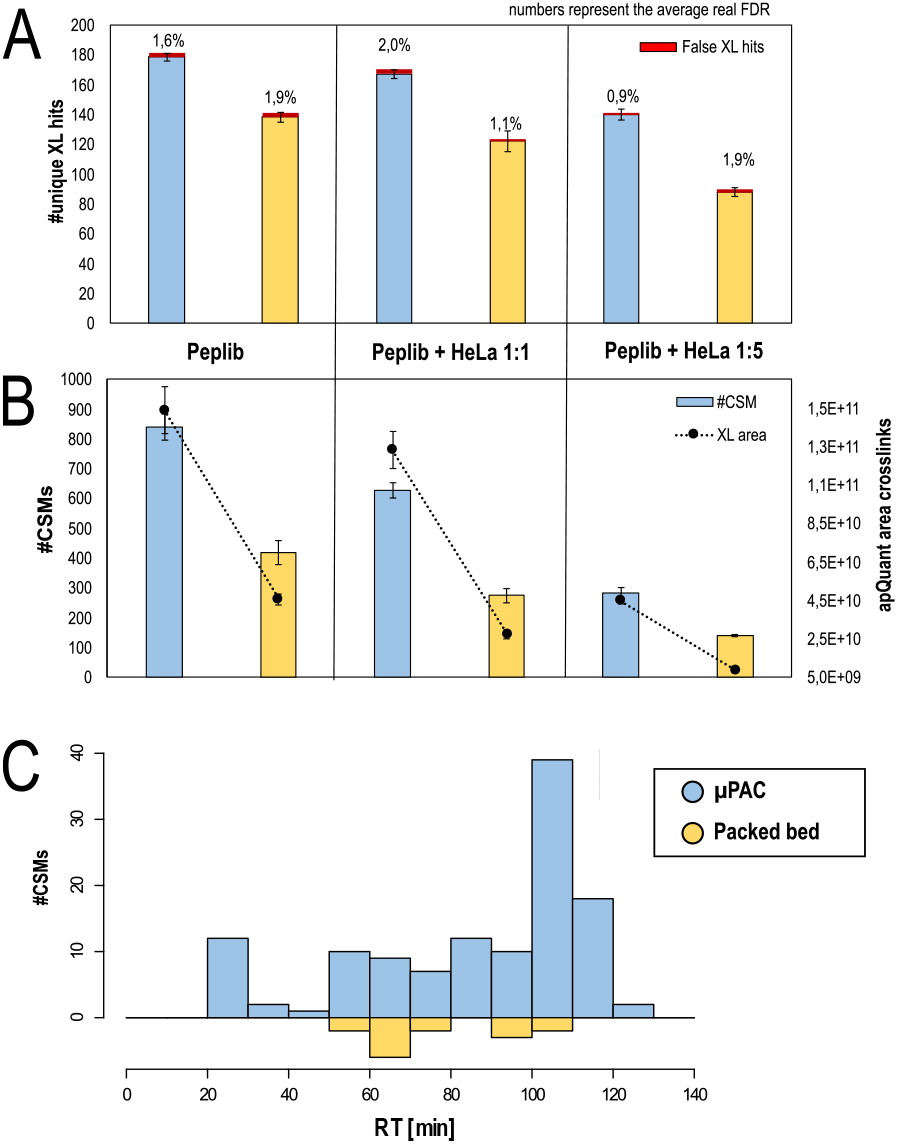
Benchmarking of generation 2 pillar array column vs packed bed column using a DSBU cross-linked synthetic peptide library. A: Number of unique cross-links identified on 1% estimated FDR level and real FDR printed above. B: B: Number of identified cross-linked peptides (CSMs) and its relative abundance based on LFQ. A and C: All values represent average values (n=3, injection replicate) with error bars depicting standard deviations. C: Number of cross-linked peptides exclusively identified with either pillar array or packed bed chromatographic setup vs retention time in on representative replicate, summed to 10 min windows.

## CONCLUSION

Data compiled in this manuscript provide a transparent perspective on the benefits that next-generation μPAC technology can bring to nanoLC-MS proteomics workflows. By combining this technology with the latest innovations in LC-MS/MS instrumentation, significant improvements in proteome coverage can be obtained with high reproducibility, robust operation and minimal sample carry over. Improvements in chromatographic performance were achieved by reducing pillar diameter and interpillar distance by a factor of 2, resulting in separation channels filled with 2.5 μm diameter pillars at an interpillar distance of 1.25 μm. As opposed to packed bed columns with integrated emitter tips, LC column and ESI emitter lifetime can be detached, providing a potentially more sustainable LC-MS solution without compromising separation performance. After optimization of a confined set of MS and LC parameters, systematically higher proteome coverage could be obtained as compared to pulled-tip packed bed nanoLC columns.

For short gradients (10 min) and limited sample amounts (10-100 ng of cell lysate digest), the impact on proteome coverage was found to be most pronounced, with gains in proteome coverage between 20 and 30% on the protein group level. When extending gradient lengths (60, 120 and 180 min) and injecting sample amounts typically encountered in the analysis of whole cell lysates (250-2000 ng), increases in coverage were less distinct, producing an increase in proteome coverage between 3 and 6% on the protein group level. The highest proteome coverage was obtained with an optimized 180 min gradient separation where 2 μg of HeLa cell digest was injected, resulting in an average number of identified protein groups of 8100.

Comparison of peptide elution behavior revealed that a larger portion of uniquely identified peptides was acquired at later eluting times, suggesting that the intrinsic difference in surface morphology (superficially porous versus fully porous) produces alternative distribution of peptides across the solvent gradient. These differences in surface morphology are also thought to be the main contributor to the reduced sample carry over that was observed in the current experiments. Column-related carry over could be reduced by a factor of 2-3 by switching from fully porous packed bed to superficially porous microfabricated column types. In-depth investigation of carry over revealed that at least 2 wash cycles were needed to wash away highly abundant peptides on traditional fully porous silica-based LC columns. Conversely, only a single wash cycle was sufficient when operating μPAC columns, hereby providing better quality data at increased instrument productivity.

Even though benchmarking studies and LC-MS/MS instrument optimization are typically performed with highly validated mammalian protein digest standards, results are often interpreted as deceiving as experiments are performed under ideal sample loading and composition conditions. To confirm results obtained in the benchmarking study, both column types were subsequently used in the analysis of a DSBU crosslinked synthetic library. When using the μPAC column, 29 to 59 more unique crosslinked peptides could be identified at a experimùentally validated FDR of 1-2%, providing additional proof for the general applicability of the next-generation μPAC technology in a range of nanoLC-MS proteomics workflows.

## SUPPORTING INFORMATION

The Supporting Information has been made available free of charge.

## AUTHOR INFORMATION

### Competing interests

JODB, AB and PJ are employees of Thermo Fisher Scientific.

## ACKNOWLEDGEMENTS

This work is supported by the EPIC-XS, Project Number 823839, funded by the Horizon 2020 Program of the European Union, by the project LS20-079 of the Vienna Science and Technology Fund and the by the ERA-CAPS I 3686 and P35045-B project of the Austrian Science Fund. We thank the IMP for general funding and access to infrastructure and especially the technicians of the protein chemistry facility for continuous laboratory support.

The authors are thankful to Christopher Pynn, Roman Huguet, and Guenther Muellauer for providing technical LC-MS support.

Spectrometry proteomics data have been deposited to the ProteomeXchange Consortium via the PRIDE 31313131 partner repository with the dataset identifier PXD030827.

Username

Password: eSyjXO16

## Notes

### Competing Interest Statement

JODB, OB and PJ are employees of Thermo Fisher Scientific

### Summary of Updates

In order to increase clarity and improve readability, we have put more emphasis on the actual results and have moved the method optimization section to the supplementary information. As a validation of the experimental results we obtained, have also included some additional testing with regards to column-to-column reproducibility.

