## Supplementary information for "Deep proteome profiling with reduced carry over using superficially porous microfabricated nanoLC columns"

### CONTENTS

**MS settings optimization ……………………………………………………………………S2**

**FAIMS setting selection……………………………………………………………………S3**

**LC conditions optimization…………………………………………………………………S3**

**Equations……………………………………………………………………………………S4**

**Figure S1……………………………………………………………………………………S5**

**Figure S2……………………………………………………………………………………S6**

**Figure S3……………………………………………………………………………………S7**

**Figure S4……………………………………………………………………………………S8**

**Figure S5……………………………………………………………………………………S9**

**Figure S6……………………………………………………………………………………S10**

**Table S1……………………………………………………………………………………S11**

**Table S2……………………………………………………………………………………S12**

**Table S3……………………………………………………………………………………S12**

**Table S4……………………………………………………………………………………S12**

**Figure S7……………………………………………………………………………………S13**

**Figure S8……………………………………………………………………………………S14**

**Figure S9……………………………………………………………………………………S15**

**Figure S10…………………………………………………………………………………S16**

Figure S11……………………………………………………………………………………………………………..……S17

Figure S12.…………………………………………………………………………………………………………………..S18

References……………………………………………………………………………………………………………………..S19

**MS settings optimization**

Previous studies have already demonstrated the benefits of performing systematic optimization of MS settings when operating orbitrap based mass analyzers in either collision induced dissociation (CID) or higher energy collision dissociation (HCD) - data dependent acquisition (DDA) modes [1–4]. The settings resulting in the highest identification rates depend to a large extent on the availability of precursor ions and therefore vary with chromatographic performance, sample loading conditions and ionization efficiency. As this is an entirely new LC-MS/MS setup combining the recently introduced Vanquish Neo UHPLC, a second-generation prototype µPAC nanoLC column and an orbitrap Eclipse Tribrid MS equipped with a FAIMS Pro interface, we deemed it was mandatory to get a proper view on the optimal MS settings for different workflow demands before starting benchmarking experiments. To cover a broad operation range, three different gradient length and sample loading combinations were defined. The effect of maximum injection time in the (linear) ion trap (MIT) and dynamic exclusion time (DET) were evaluated while keeping all other MS and LC settings constant (Figure S3 – Table S1). Short gradients produce sharper peaks and generate higher relative detection responses, which reduces the amount of sample material needed to continuously trigger MS/MS events. As no significant impact was anticipated by loading micrograms of sample material when analysis time is limited, a short method with relatively low sample loads (10 min gradient / 50 ng of HeLa cell digest) was used to optimize settings for high throughput analyses where high sensitivity is needed. Separation performance (peak capacity) can be increased by extending the LC solvent gradient, but this will result in a reduction of the relative concentration at which peptides elute[5–7]. As a result, the concentration of low abundant peptides will drop below the limit needed to trigger MS/MS and more material must be loaded to convert increased LC separation performance into an increase in ID's. A routine method with standard sample loads (60 min gradient / 1 µg of HeLa cell digest) was used to cover typical nanoLC-MS bottom-up proteomics conditions and a long method with high sample loads (180 min gradient / 3 µg of HeLa cell digest) was used to explore deeper proteome coverage.

To get the highest possible scan speed, the instrument was operated in high-low mode where the ion trap (IT) rather than the orbitrap (OT) is used for MS2 spectrum acquisition. Theoretically, operating the Orbitrap Eclipse in OT-IT mode with ion trap speed set at Turbo rate and covering a scan width of 1200 m/z allows an approximate MS scanning rate of 45 Hz [8–10]. Due to time spent for MS1 scans we could achieve MS2 scan rates of up to 35 Hz with our DDA methods, and this for all three conditions tested during LC-MS/MS optimization (Figure S4). The highest scan rates were consistently achieved at low MIT values, where ion accumulation times do not exceed MS/MS scan duration and associated overhead time. A stable MS2 scan speed of 35 Hz is observed up to MIT of 15 ms, which is in line with earlier reports where optimal MIT settings have been evaluated for a range of m/z ranges and ion trap speed settings[8]. MIT of approximately 15 ms also appeared to be the sweet spot in our analyses (Figure S3 A-C). When MIT is increased above 15 ms, a linear decline in MS2 scanning speed with concomitant decrease in absolute identification numbers is observed. This highlights the importance of MS scan speed towards maximizing feature detection. Even though MS2 scanning speed was improved further, injection times below 15 ms did not result in higher MS2 identification rates nor absolute identification numbers. By collecting fewer ions for MS/MS, overall spectral quality deteriorates. Optimal MIT settings appear to be near the inflection point where the MS is still scanning as fast as possible but producing MS spectra with the highest possible quality.

DET is another indispensable setting in DDA acquisition mode. It enables the rejection of high abundant (and often 'broader' eluting) peptides from redundant sampling and fragmentation. Redundant sampling typically results in higher absolute PSM numbers but lower overall peptide and protein identifications. Time spent on the fragmentation of a peptide that has been already sampled reduces the time that can be spent to find new ones. This is observed when low DET values were evaluated. The optimal DET setting typically depends on the elution width of high to medium abundant peptides impacted by the LC column performance and gradient length[11]. Even though we found that DET settings only affect MS scanning rates to a very limited extent, a significant impact on absolute identification numbers was observed (Figure S3 D-F). The optimum DET setting is directly related to the observed peak width and therefore can be gauged by plotting peak width distributions for a certain separation condition. Below the optimum DET value, identifications are lost due to redundant sampling. Above the optimum, identifications are lost because the MS instrument is running out of precursors and not using up all available speed.

Even though the absolute sample load is significantly different, the concentration distribution at which peptides are presented to the MS is not that divergent. With median PSM intensities between 0.5 and 1.5 x $10^6$ (Figure S8), optimal MIT settings were found to be quite similar for all three LC methods. "PSM intensity" is the intensity of the precursor in the MS1 scan preceding the MS2 scan. This intensity is reported in charges per second to account for differences in MS1 fill time. As expected, optimal DET settings do however shift with increased peak width, leading to higher optimal values for longer gradients and higher sample loads. Based on the comprehensive results obtained during this optimization, we defined a set of optimal MS settings for the subsequent benchmarking experiment (Table S4).

### FAIMS settings selection

In accordance with previous reports on using FAIMS Pro in internal compensation voltage (CV) stepping mode, a 3 CV internal stepping method (-45, -55 and -75) with CVs that are 10–20 V apart and centered near the identification distribution maximum was used to evaluate the proteome coverage for different LC methods[12–14]. It was beyond the scope of the current study to provide an in-depth evaluation of the effect of different compensation voltages or to quantify the additional depth that can be generated by implementing FAIMS-MS. The results obtained for initial column installation runs (QC gradient of 15 min) quickly pointed out the added value of using more CV's when aiming at comprehensive proteome coverage, 23% more protein groups (3002 vs 2431) could be identified from 100 ng of HeLa cell digest when increasing CV steps from 2 to 3. For LC gradient lengths up to 120 min, we did not evaluate alternative FAIMS settings, all experiments were performed using a 3CV method that cycles between CVs at an interval of 1s, generating one MS1 scan for each FAIMS CV every 3s. For the longest LC gradient tested (180 min), the potential of an additional compensation voltage was however evaluated as broader peptide elution allows more time available per each CV. Proteome coverage obtained for the 3 CV method was compared to what could be obtained with 4 different CV values (Figure S5 – Table S2). Without claiming that this is the unique optimal combination of CV values, we obtained maximum proteome coverage when stepping between CV values of -45, -55, -65 and -75V.

#### LC conditions optimization

As from the first reports on the characteristics of nanoelectrospray, great potential has been anticipated for the combination of low liquid flow rates with ESI-MS[15]. As current supplied by electrospray is proportional to the square root of the flow rate, liquid is ejected by electrospray at higher charge densities at low flow rates[16]. This typically results in higher ionization efficiency of analytes and improves detection sensitivity. Increases in detection sensitivity have proven to be of key importance in the pursuit of comprehensive proteome characterization. Improvements in the field have typically been achieved by increasing the depth to which precursor molecules could be sampled. In search of improved detection sensitivity for limited sample amounts, the implementation of ultra-low flow (ULF) ESI-MS has seen quite a revival in the last few years. Major progress has been documented when ULF (flow rates below 100 nL/min) was combined with ESI-MS [17–22]. Robust and routine operation at these low flow rates is however not straightforward and often required highly specialized LC systems or customized pre-column flow splitting configurations. The importance of accurate flow rate control and gradient formation precision cannot be underestimated for practical implementations as these parameters define quantitation accuracy to a large extent. Operating columns at low flow rates typically comes at the cost of increased analysis and overhead time. To restrict the impact on total analysis time and at the same time ensure good chromatographic performance, it is crucial to align LC column dimensions with the desired flow rate range. The prototype μPAC column has a reduced cross-section (equivalent to a packed bed column with ID of 60 μm) and a total column volume of approximately 1,5 μL. This holds great potential for operation at flow rates lower than 300 nL/min. To find the sweet spot for comprehensive proteome analysis, we evaluated the effect of decreased LC flow rate on chromatographic performance and proteome coverage. The recently introduced Vanquish Neo UHPLC system allows setting the flow from 1 nL/min up to 100 μL/min with 1 nL/min increments and running gradients at typical nano/cap/micro LC flow rates as well as ULF without flow splitting or hardware changes. This is enabled by active flow control and multipoint flow calibration that do not require re-adjustment or re-calibration during the LC usage. The wide flow range LC capabilities and low gradient delay volume together with system operation at constant maximum pressure specified for the column (400 bar) during the sample loading and column equilibration reduced the overhead time and made it possible to run gradients starting from 50 nL/min. Keeping all settings other than flow rate constant, we systematically compared the metrics obtained for a 180 min gradient separation (1 μg HeLa digest sample on column). The gradient was identical for all flow rates tested and has been described in the experimental section. MS acquisition times were adapted to compensate increasing void times at lower flow rates. Base peak chromatograms obtained at flow rates ranging from 50 to 300 nL/min demonstrate the impact of flow rate on the peptide elution start (Figure S6 C-H). Whereas the first peptides elute after approximately 6 min at 300 nL/min, operation at 50 nL/min postpones elution by a factor of 6. Even though the actual elution window in which digested protein material elutes was similar for all flow rates tested (180 min), a significant increase in absolute signal intensity was observed when reducing flow rate. This was confirmed when comparing mean PSM intensities obtained at different flow rates. Surprisingly, the increase in ionization efficiency did not result in improved proteome coverage. On the contrary, optimal proteome coverage (protein group IDs) was obtained at a flow rate of approximately 200 nL/min where a compromise between chromatographic performance (FWHM in figure S6) and ionization efficiency (mean PSM intensity in figure S6) was achieved. The increased ionization efficiency observed at low flow rates does however suggest that another cycle of MS settings optimization might have been needed to explore the full potential of ULF LC ESI MS on this column. We did not pursue low flow rate operation any further as the current evaluation was aimed at finding a balance between MS utilization time, sensitivity for typical bottom-up proteomics sample loads (250 - 2000 ng), and separation quality. LC-MS settings used during flow rate optimization are listed in Table S3.

### Equations

**FWHM based Peak capacity** (Equation 1)

$$n_{C\ 50\%} = \left(\frac{T_{Gradient}}{median\ FWHM}\right) + 1$$

**Instrument productivity** (Equation 2)

$$Instrument\ productivity = \frac{Effective\ peptide\ elution\ window}{Instrument\ method\ duration + overhead\ time}$$

**Relative percentage of carry-over** (Equation 3)

$$Relative\ \%\ carry\ over = \frac{Number\ of\ peptide\ groups\ identified\ in\ blank\ run}{Number\ of\ peptide\ groups\ identified\ in\ analytical\ run\ preceding\ blank\ run}$$

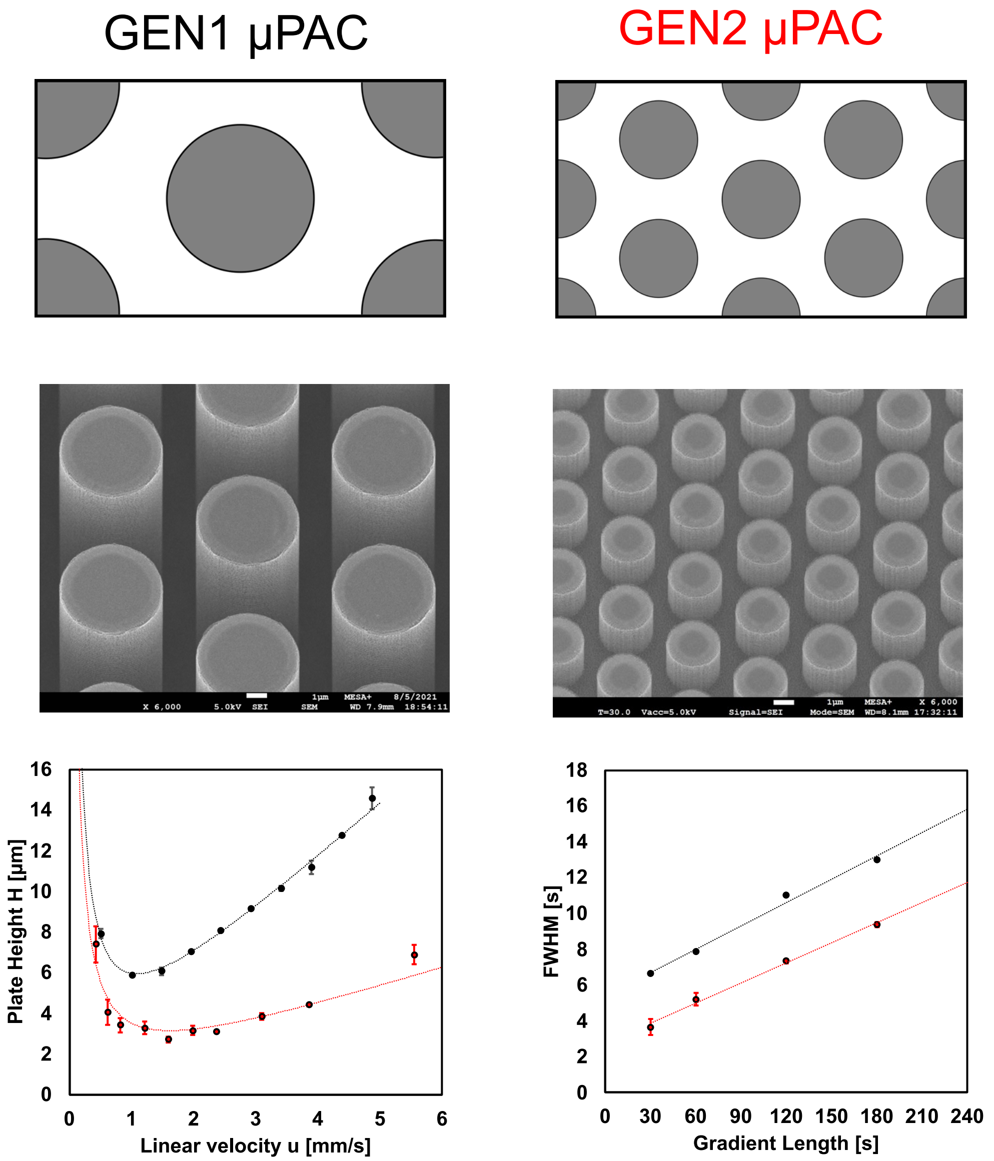


**Figure S1. Top: Schematic representation of the unit cells used to design pillar array column generations. Left: Generation 1, Right: generation 2. Middle: Scanning Electron Microscopy (SEM) images showing a birds eye view of micro pillar array stationary phase support generations. Left: generation 1, Right: generation 2. Bottom: Comparison of chromatographic performance between generations in either isocratic (left) mode or gradient mode (right), generation 1 = black, generation 2 = red.**

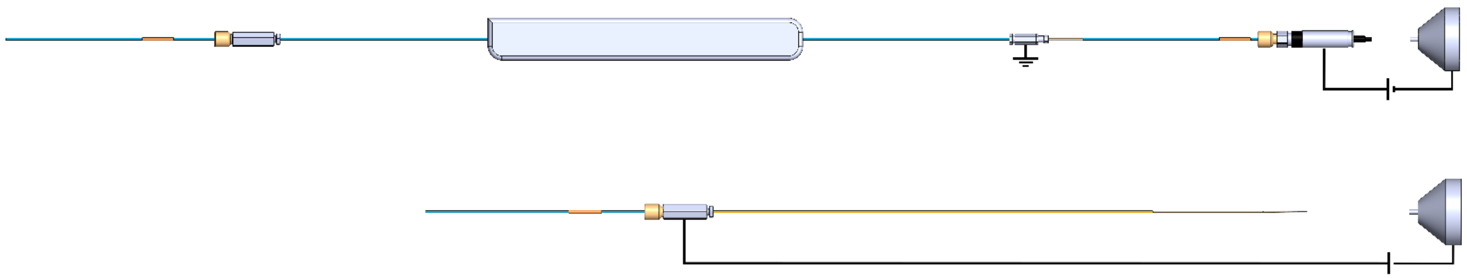

**Figure S2. Schematic overview of the fluidic and electrical configurations used to operate packed bed pulled tip emitter type columns (Bottom) and prototype µPAC columns (Top).**

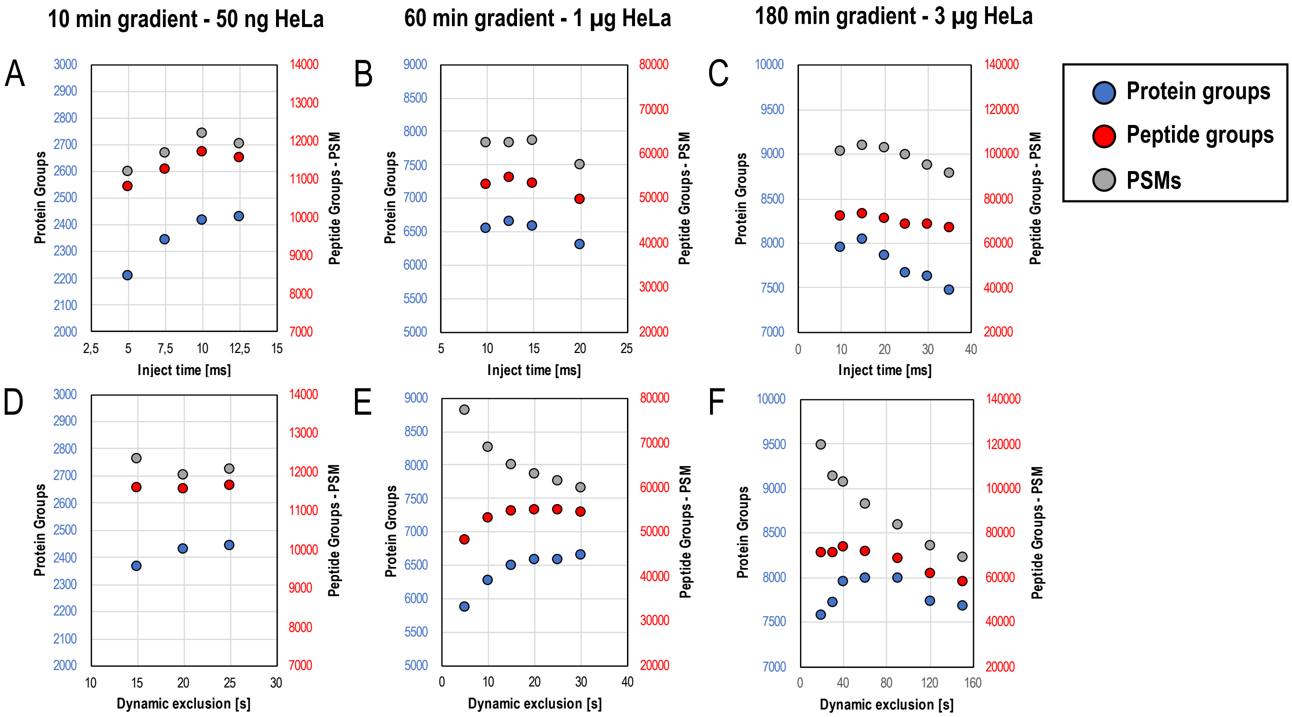


**Figure S3: Impact of MS settings on identification rate – throughput (10 min gradient) versus routine (60 min gradient) and comprehensive (180 min gradient) (A-C): Impact of maximum injection time (MIT) in the ion trap, (D-F): Impact of dynamic exclusion time (DET). Blue = Protein Groups, Red = Peptide Groups, Grey = PSMs. 50 cm prototype µPAC column was used.**

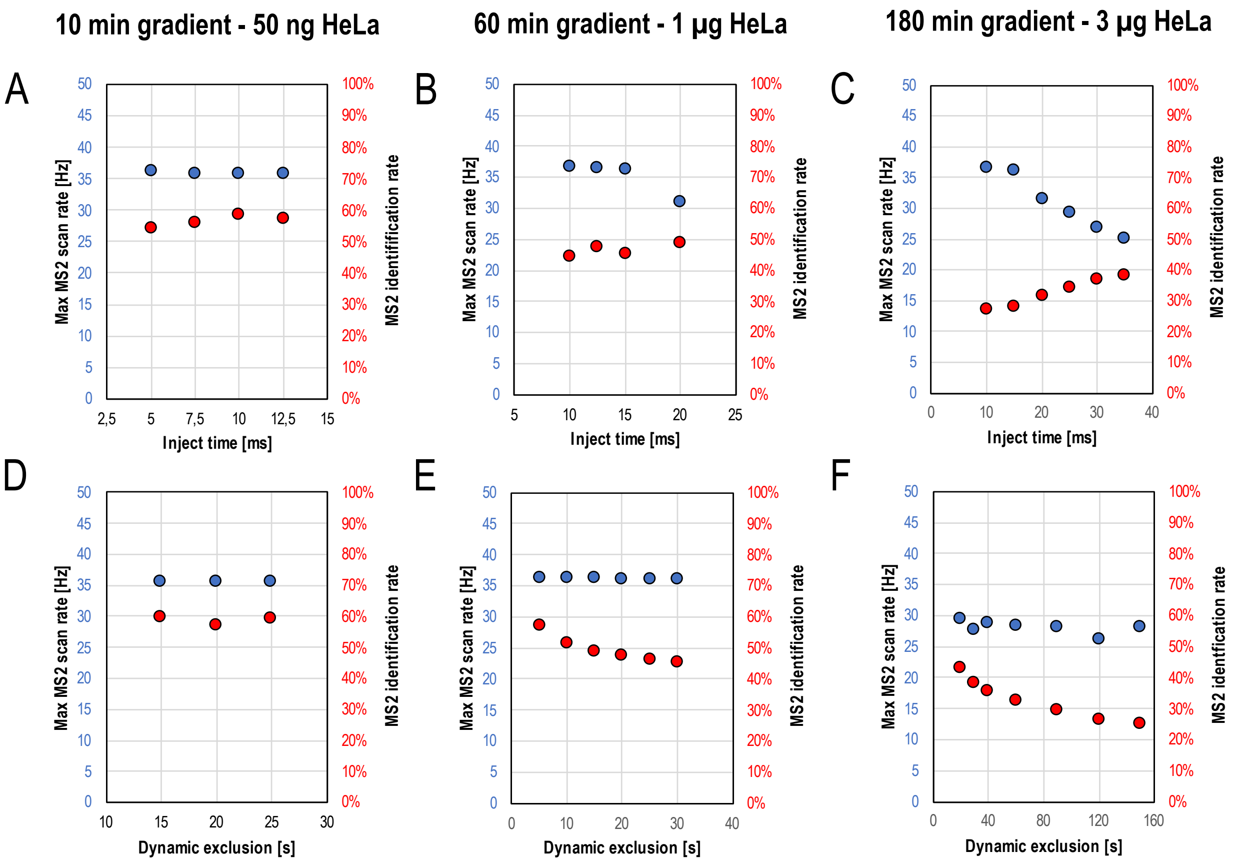


**Figure S4. Impact of MS settings on MS scanning speed and MS2 identification rate – throughput (10 min gradient) versus routine (60 min gradient) and comprehensive (180 min gradient) (A-C): Impact of maximum injection time (MIT) in the ion trap, (D-F): Impact of dynamic exclusion time (DET). Blue = max MS2 scan rate in Hz, Red = MS2 identification rate. 50 cm prototype µPAC column was used.**

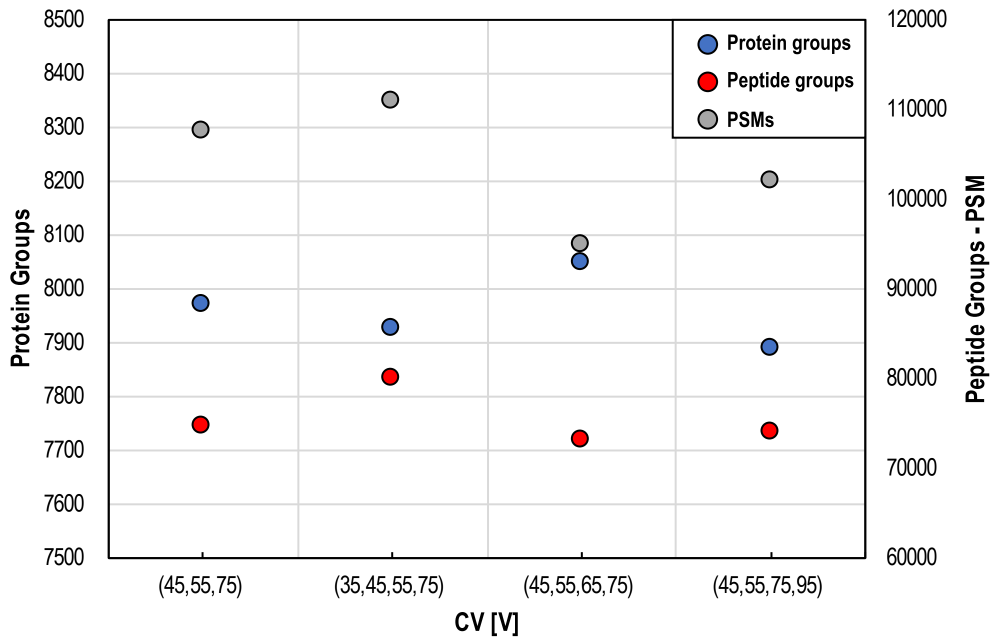


**Figure S5. Impact of FAIMS pro compensation voltage settings on proteome coverage. 1µg HeLa digest – 180 min gradient. Blue = Protein Groups, Red = Peptide Groups, Grey = PSM. 50 cm prototype µPAC column was used.**

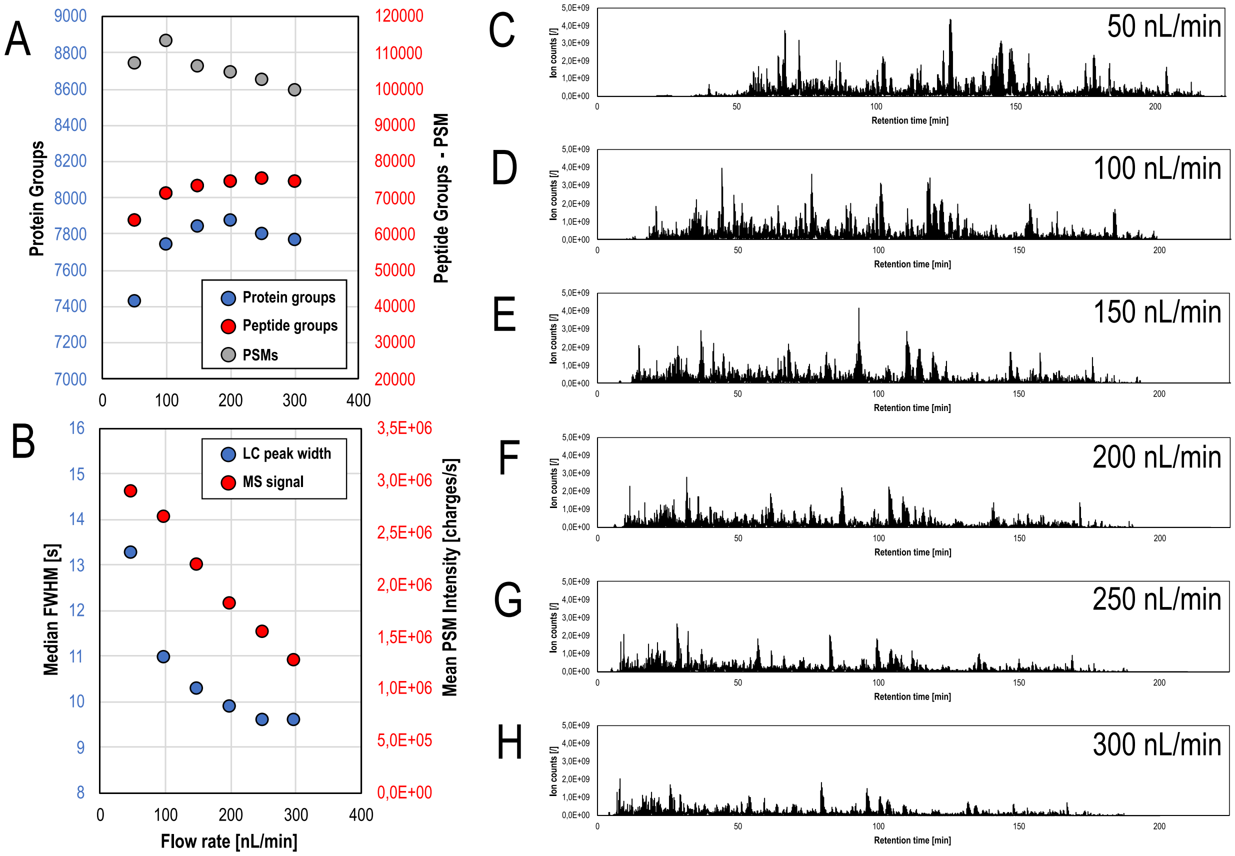


**Figure S6: Impact of flow rate on identification rate, chromatographic performance, elution profile and signal intensity. (A): ID's as a function of flow rate, Blue = Protein Groups, Red = Peptide Groups, Grey = PSMs., (B): FWHM (Blue) and Mean PSM intensity (Red) as a function of flow rate, (C-H): Base Peak Chromatogram comparison / elution window for different flow rates. 50 cm prototype µPAC column was used.**

| Gradient time [min] | LC Flow rate [nL/min] | FAIMS Compensation voltages [V] | Total MS cycle time [s] | Maximum inject time ddMS$^2$ IT [ms] | Dynamic exclusion time [s] |
|---|---|---|---|---|---|
| 10 | 300 | -45 / -55 / -75 | 3 | 5 | 20 |
| 10 | 300 | -45 / -55 / -75 | 3 | 7.5 | 20 |
| 10 | 300 | -45 / -55 / -75 | 3 | 10 | 20 |
| 10 | 300 | -45 / -55 / -75 | 3 | 12.5 | 20 |
| 10 | 300 | -45 / -55 / -75 | 3 | 12.5 | 15 |
| 10 | 300 | -45 / -55 / -75 | 3 | 12.5 | 20 |
| 10 | 300 | -45 / -55 / -75 | 3 | 12.5 | 25 |
| 60 | 300 | -45 / -55 / -75 | 3 | 10 | 20 |
| 60 | 300 | -45 / -55 / -75 | 3 | 12.5 | 20 |
| 60 | 300 | -45 / -55 / -75 | 3 | 15 | 20 |
| 60 | 300 | -45 / -55 / -75 | 3 | 20 | 20 |
| 60 | 300 | -45 / -55 / -75 | 3 | 15 | 5 |
| 60 | 300 | -45 / -55 / -75 | 3 | 15 | 10 |
| 60 | 300 | -45 / -55 / -75 | 3 | 15 | 15 |
| 60 | 300 | -45 / -55 / -75 | 3 | 15 | 20 |
| 60 | 300 | -45 / -55 / -75 | 3 | 15 | 25 |
| 60 | 300 | -45 / -55 / -75 | 3 | 15 | 30 |
| 180 | 300 | -45 / -55 / -75 | 3 | 10 | 40 |
| 180 | 300 | -45 / -55 / -75 | 3 | 15 | 40 |
| 180 | 300 | -45 / -55 / -75 | 3 | 20 | 40 |
| 180 | 300 | -45 / -55 / -75 | 3 | 25 | 40 |
| 180 | 300 | -45 / -55 / -75 | 3 | 30 | 40 |
| 180 | 300 | -45 / -55 / -75 | 3 | 35 | 40 |
| 180 | 300 | -45 / -55 / -75 | 3 | 25 | 20 |
| 180 | 300 | -45 / -55 / -75 | 3 | 25 | 30 |
| 180 | 300 | -45 / -55 / -75 | 3 | 25 | 40 |
| 180 | 300 | -45 / -55 / -75 | 3 | 25 | 60 |
| 180 | 300 | -45 / -55 / -75 | 3 | 25 | 90 |
| 180 | 300 | -45 / -55 / -75 | 3 | 25 | 120 |
| 180 | 300 | -45 / -55 / -75 | 3 | 25 | 150 |

**Table S1: LC/MS settings used during MS method optimization.**

| Gradient time [min] | LC Flow rate [nL/min] | FAIMS Compensation voltages [V] | Total MS cycle time [s] | Maximum inject time ddMS$^2$ IT [ms] | Dynamic exclusion time [s] |
|---|---|---|---|---|---|
| 180 | 200 | -45 / -55 /-75 | 3 | 15 | 20 |
| 180 | 200 | -35 / -45 / -55 / -75 | 3 | 15 | 20 |
| 180 | 200 | -45 / -55 / -65 /-75 | 3 | 15 | 20 |
| 180 | 200 | -45 / -55 / -75 /-95 | 3 | 15 | 20 |

**Table S2: LC/MS settings used during FAIMS optimization.**

| Gradient time [min] | LC Flow rate [nL/min] | FAIMS Compensation voltages [V] | Total MS cycle time [s] | Maximum inject time ddMS$^2$ IT [ms] | Dynamic exclusion time [s] |
|---|---|---|---|---|---|
| 180 | 50 | -45 / -55 / -75 | 3 | 15 | 20 |
| 180 | 100 | -45 / -55 / -75 | 3 | 15 | 20 |
| 180 | 150 | -45 / -55 / -75 | 3 | 15 | 20 |
| 180 | 200 | -45 / -55 / -75 | 3 | 15 | 20 |
| 180 | 250 | -45 / -55 / -75 | 3 | 15 | 20 |
| 180 | 300 | -45 / -55 / -75 | 3 | 15 | 20 |

**Table S3: LC/MS settings used during LC flow rate optimization.**

| Gradient time [min] | LC Flow rate [nL/min] | FAIMS Compensation voltages [V] | Total MS cycle time [s] | Maximum inject time ddMS$^2$ IT [ms] | Dynamic exclusion time [s] |
|---|---|---|---|---|---|
| 10 | 200 | -45 / -55 / -75 | 3 | 12.5 | 20 |
| 60 | 200 | -45 / -55 / -75 | 3 | 12.5 | 25 |
| 120 | 200 | -45 / -55 / -75 | 3 | 15 | 40 |
| 180 | 200 | -45 / -55 / -65 / -75 | 4 | 15 | 60 |

**Table S4: LC/MS settings used for the final benchmarking experiment.**

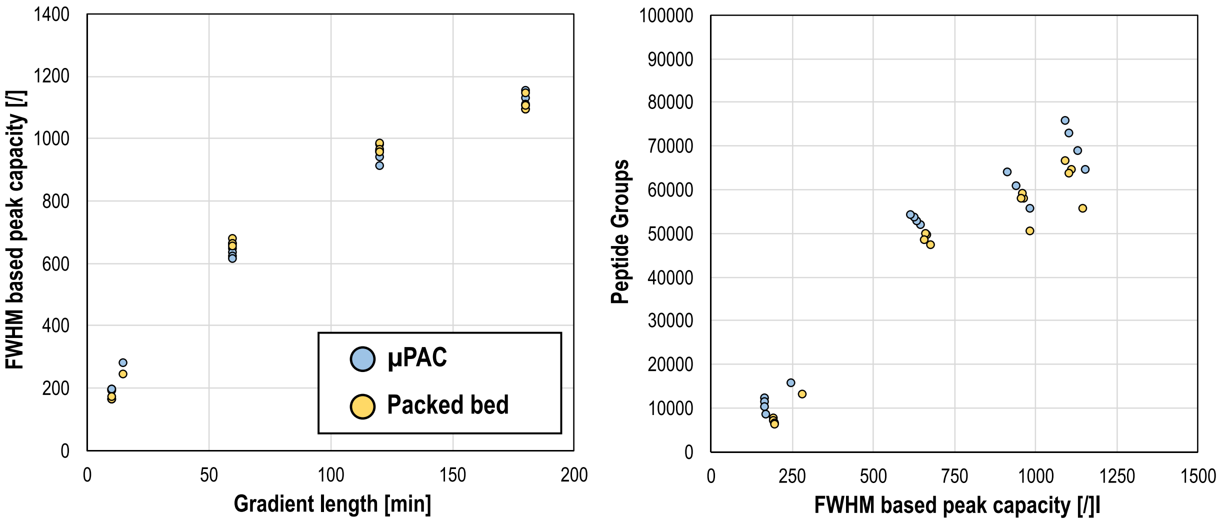


**Figure S7. FWHM based peak capacity for different column types. Peak capacity is plotted as a function of gradient time for both columns and all sample loads (A). Number of Identified peptide groups plotted as a function of peak capacity, both column types, all sample loads (10-2000 ng HeLa) (B).**

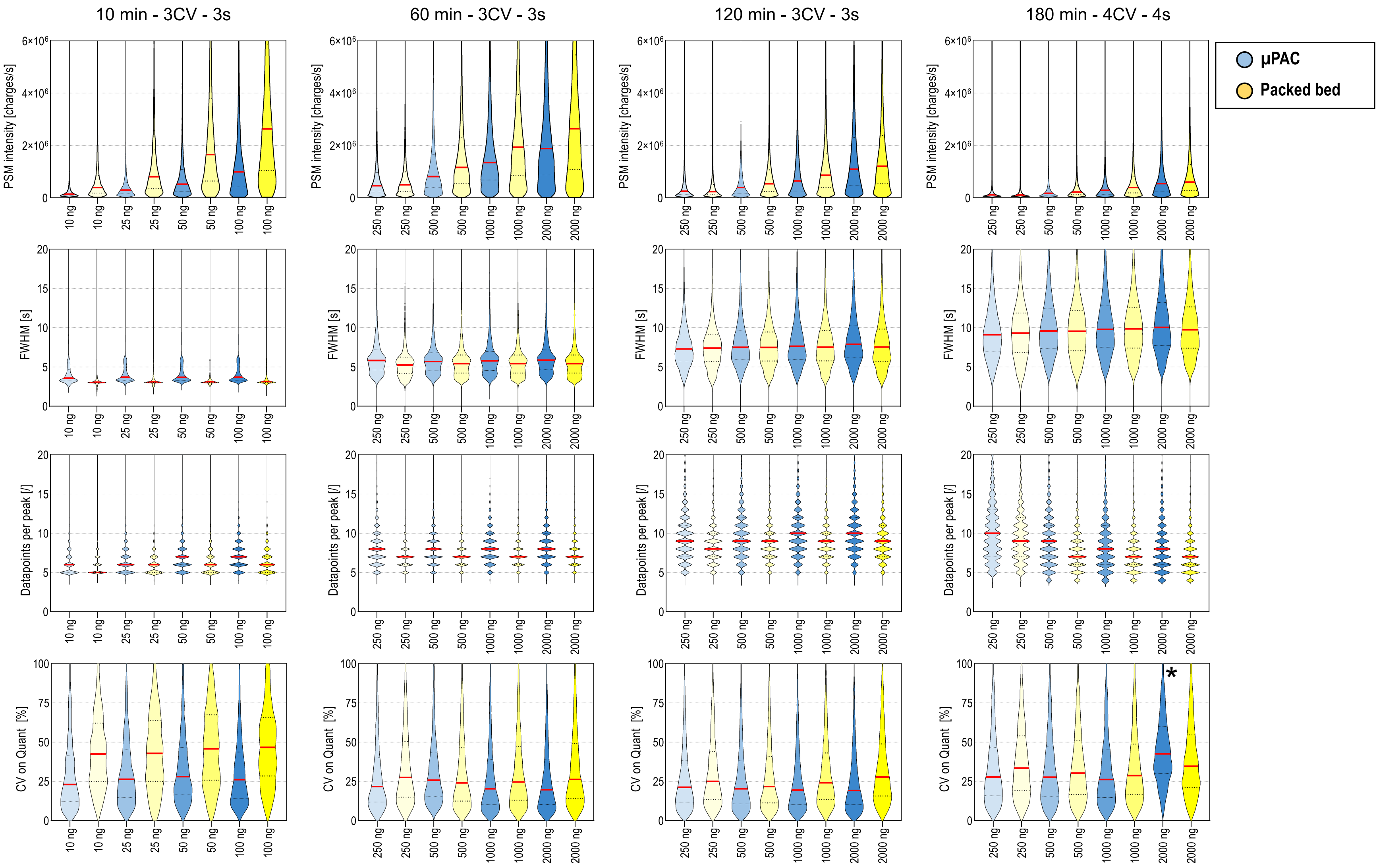


**Figure S8. Violin plots showing the distribution (from top to bottom) of PSM intensity, FWHM, Datapoints per peak and %CV on quantification (n=3). Conditions used during the final benchmarking experiment are shown. Median values shown in red. * CV for the 2000 ng sample 180 min analysis was higher due to inclusion of an outlier.**

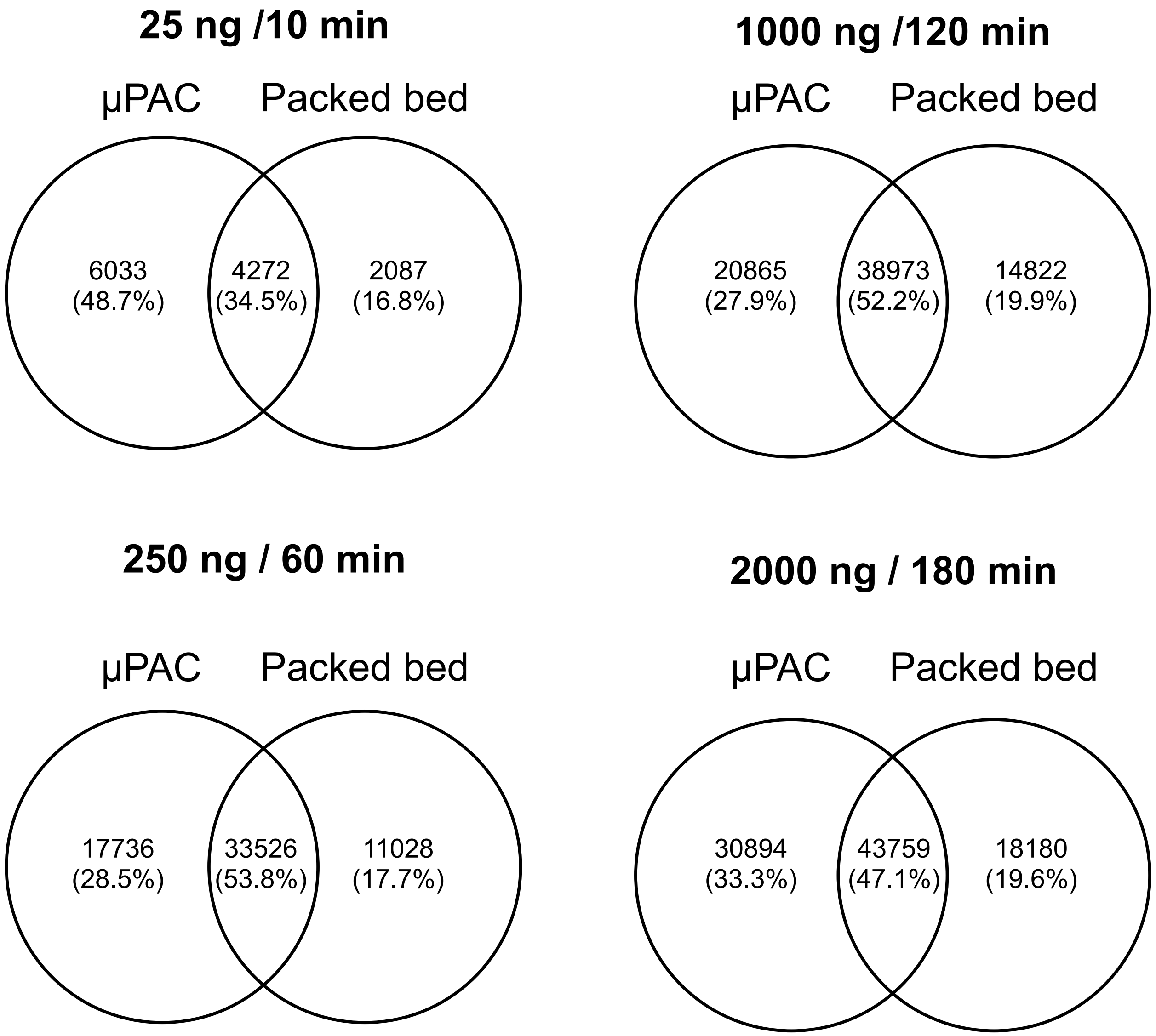


**Figure S9. Venn diagrams showing the overlap between identified peptide groups on the generation 2 µPAC column and the packed bed column.**

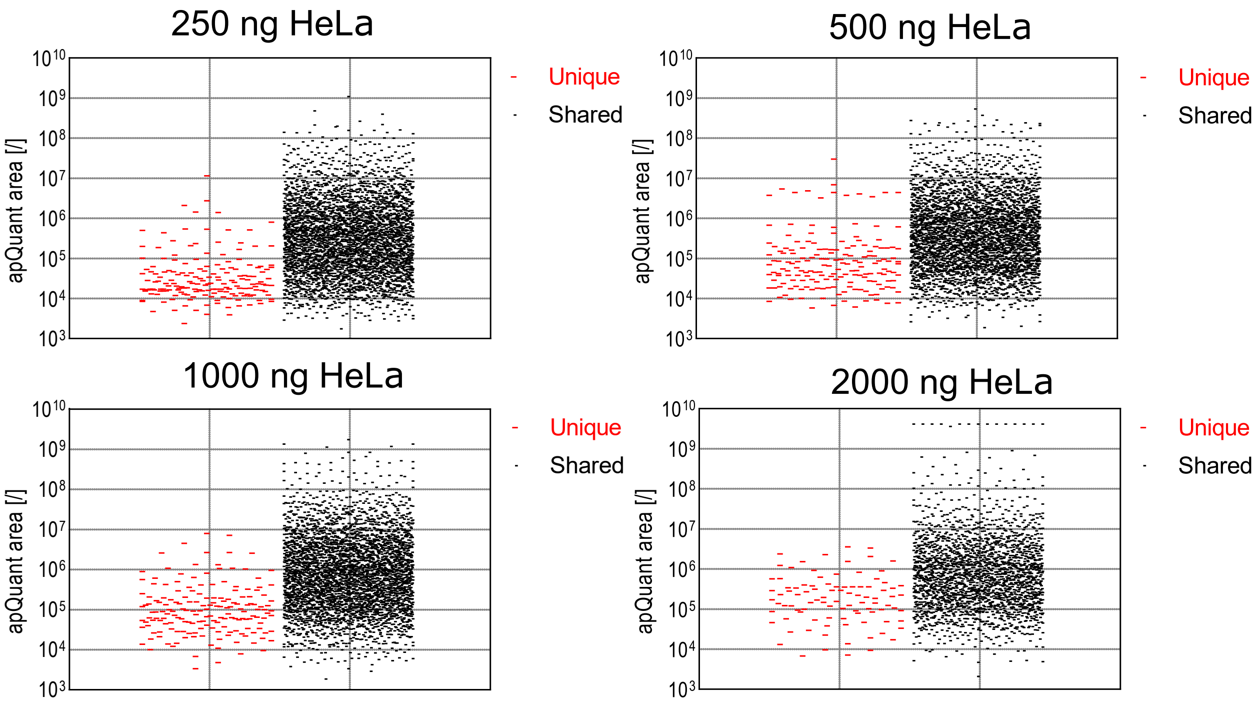


**Figure S10. Comparison of apQuant area obtained for peptides that were either uniquely identified in the 180 min gradient method or shared with the 60 and 120 min gradient method. 50 cm prototype µPAC column was used.**

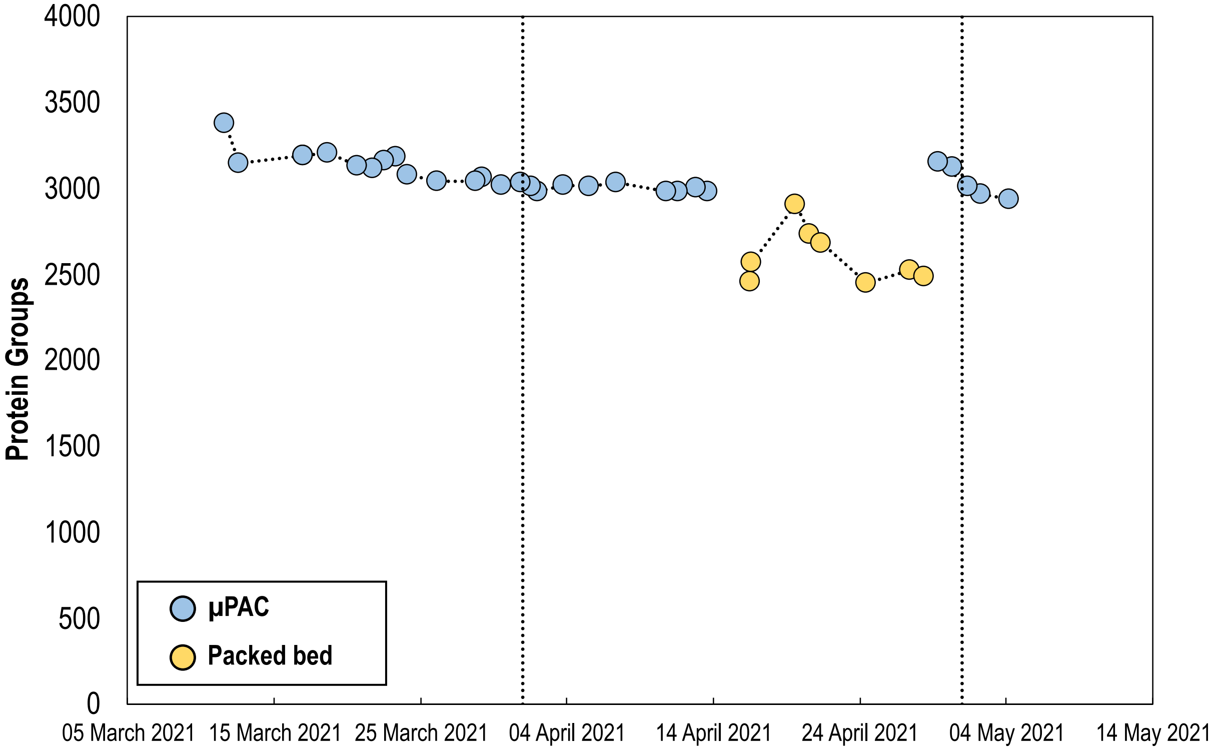


**Figure S11. Evolution of column performance (Protein Group ID's) for 15 min gradient QC runs over time. 100 ng HeLa. Blue = GEN2 µPAC (n=2), Yellow = packed bed (n=1)**

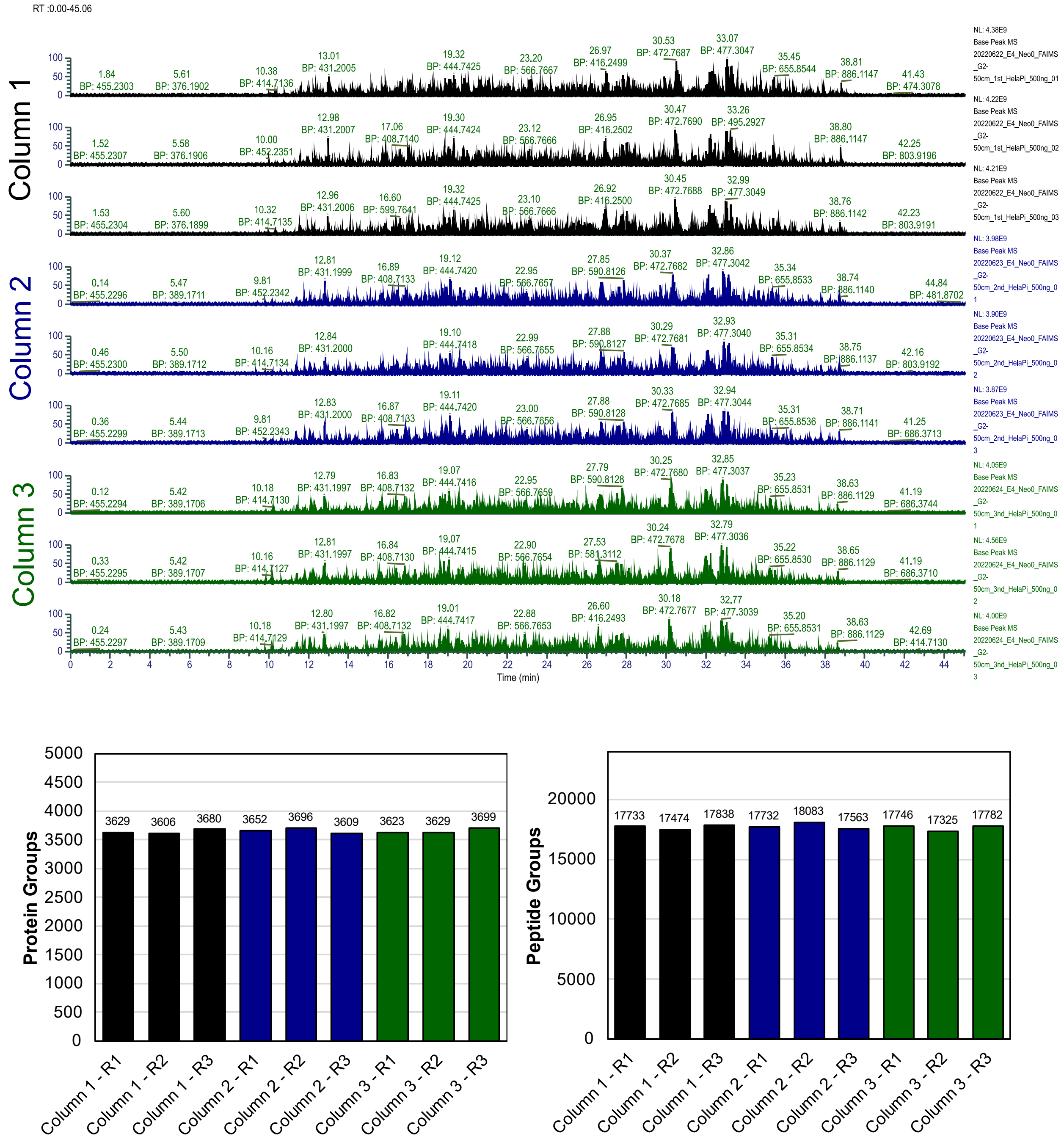


**Figure S12. Column reproducibility evaluation on an Orbitrap Exploris 480 instrument using a similar LC instrument (Vanquish Neo UHPLC) as used in to generate data published in this contribution. Top: basepeak chromatograms obtained for 30 min gradient separation of 500 ng HeLa digest on different generation 2 µPAC columns. (Columns, n=3 - injection replicates, n=3). Bottom: Protein and peptide group identifications obtained.**
